# Contrasting amphibian population trend from a protected area in Madagascar reveal severe underestimation of extinction risks

**DOI:** 10.1101/2025.03.28.645942

**Authors:** Xavier Porcel, Angelica Crottini, Karen Freeman, Jean Noёl, Jean Honoré Velo, Honoré Lava, Georges Rendrirendry, Franco Andreone, Nicolas Dubos

## Abstract

The amphibian endemic fauna from Madagascar is currently facing multiple threats, including habitat loss, climate change and invasive species. Many species are listed as threatened by the IUCN, but long-term monitoring is scarce and assessments rarely account for population trends. Using a 13-year amphibian survey in the Strict Nature Reserve of Betampona (eastern Madagascar), we assessed the population trends of 38 species between 2010 and 2022 based on amphibian activity. Despite the high protecction status of the reserve, we found a negative trend for 15 species, a positive trend for 13 species increasing and 13 species with no significant trend detected. We propose to update the status of 15 species towards a threatened category (CR, EN, VU). Contrasting population trends in amphibians from Betampona might modify species composition and ecosystem functions in the future. This study highlights an important underestimation of extinction risks for a large proportion of amphibians from Madagascar, and more generally in the tropics where long-term population trends are poorly documented.

## I – Introduction

The World’s biodiversity is currently suffering from a massive global extinction crisis, qualified by some authors as the 6th mass extinction (Barnosky et al., 2011; Ceballos and Ehrlich, 2010). Among all the taxa, amphibians are the most threatened, ahead of birds and mammals, with about 3000 species listed as Threatened (UICN 2024). The decline of amphibians sometimes remains enigmatic (Blaustein and Kiesecker, 2002; Collins et al., 2003; Gibbon et al., 2000), but major causes identified are the destruction and fragmentation of their habitats, climate change and disease (Cayuela et aL Submitted). Among the other causes of the decline, amphibian suffer from introduced invasive species, environmental pollution, parasitism and unsustainable use (Andreone et al., 2021; Collins et al., 2003; Cunningham, 2018; Gibbon et al., 2000; Meredith et al., 2016). The global decline of amphibian results probably from a synergy between all of these factors (Blaustein and Kiesecker, 2002; Collins et al., 2003; Meredith et al., 2016). Notwithstanding the urgent need for conservation action, an important proportion of species remain unassessed, especially in the tropics (Caetano et al., 2022). Conservation assessments can rely on estimations of population trends (i.e. criterion A; IUCN Standards and Petitions Committee 2024) (Mancini et al., 2024). However, long-term monitoring is rarely available and there is a huge lack of information on species temporal trends and species turnover on amphibian communities.

Amphibian populations and communities dwelling in forests are highly exposed to habitat fragmentation (Vallan, 2000). That is mainly because amphibians are poor dispersers and forest species are highly dependent on habitat specificities (e.g. litter, canopy cover; (Todd et al., 2009)). In Madagascar, the loss of forest habitats is predicted to continue and increase in the coming years (Vieilledent et al., 2018).

Madagascar is a large country with almost unrivaled levels of endemism, considered as a hotspot of amphibian and global biodiversity (Myers et al. 2000, The Global Amphibian Assessment (GAA) 2005). The latest complete inventory led to the listing of 292 nominal species (Perl et al., 2014; Vieites et al., 2009) and identify several “candidate species” defined by individuals whose genetic sequences show divergences from others above a certain defined threshold (Vences and Wake, 2007). Authors consider that thanks to DNA progress, the number of Malagasy Anura species could reach more than 530 species (Perl et al., 2014). Considering the endemicity, 100 % of the native species are strictly endemic to Madagascar and its inshore islands, making this island an incredibly rich place where conservation effort needs to be focus on.

The highest amphibian diversity and abundance in Madagascar is found along the oriental coast, in the eastern rainforest corridor (Brown et al., 2016) where deforestation has been extensive (Vieilledent et al., 2018). The last remnant forests are isolated patches, often very small, but may contain a large number of amphibian species, some of which are often micro-endemic to these forests and to specific conditions inside them (Rosa et al., 2012). However, because of their small size, these fragments are probably subject to an important edge effect as well as Allee effect, therefore highly exposed to changes in both biotic and abiotic parameters of the forests (Bourgoin et al., 2024; Slater et al., 2024). Abrupt climatic events triggered by global change could also have a major impact, promoting the emergence of invasive exotic species (Dubos et al., 2023), leading to the homogenization of the environment (Clavel et al., 2011) and the loss of niches, especially those of specialist species (Dubos et al., 2022). The Strict Nature Reserve of Betampona is one of these forest fragments.

In Madagascar, and more generally in the tropics, long-term monitoring is rarely available and there is a huge lack of information on species temporal trends and species turnover on amphibian communities. Relying on 13 years of monitoring from the Betampona Strict Nature Reserve, we assessed the temporal trends in amphibian activity at the community and population levels. Based on these trends, we classified the trends according to the IUCN criteria A4 (ongoing population size reduction) and provide recommendations for species conservation status. We further discuss the potential consequences of population trends on species composition and ecosystem functions.

## II – Material & Methods

### II.1 – Study site

Betampona forest (E49°12’00”–49°15’00”, S17°15’00”–17°55’00”) (**Figure 1**) is an isolated 2,228-hectare lowland rainforest fragment located approximately 40km northwest of Toamasina. This forest extends from 92 to 571 m above sea level and is characterized by a tropical rainforest climate. The area comprises water sources like fast and slow-moving streams and hills (Rosa et al., 2012). This forest became the first area to receive the status of Strict Nature Reserve (“*Réserve Naturelle Intégrale*”) in 1927 and is now managed by the Madagascar National Parks (MNP) NGO. The locally based Madagascar Fauna and Flora Group (MFG) is MNP’s official research partner for the reserve and is promoting the conservation and scientific study of this ecosystem, leading to an increasing scientific interest over the past decade (Andreone et al., 2010; Bellati et al., 2018; Dubos et al., 2020; Rosa et al., 2014, 2012, 2011). More recent surveys have revised Betampona species’ list that now includes over 80 frog species (A. Crottini, pers. obs.). Based on previous inventory efforts in Betampona, we selected two amphibian-rich sites to establish long-term monitoring transects for amphibians, locally known as “Sahabefoza” (S 17° 54′ 51.2″; E 049° 12′ 27.7″) and “Sahambendrana” (S 17° 53′ 54.2″; E 049° 12′ 55.4″) (**Figure 1**).

**Figure 1:**
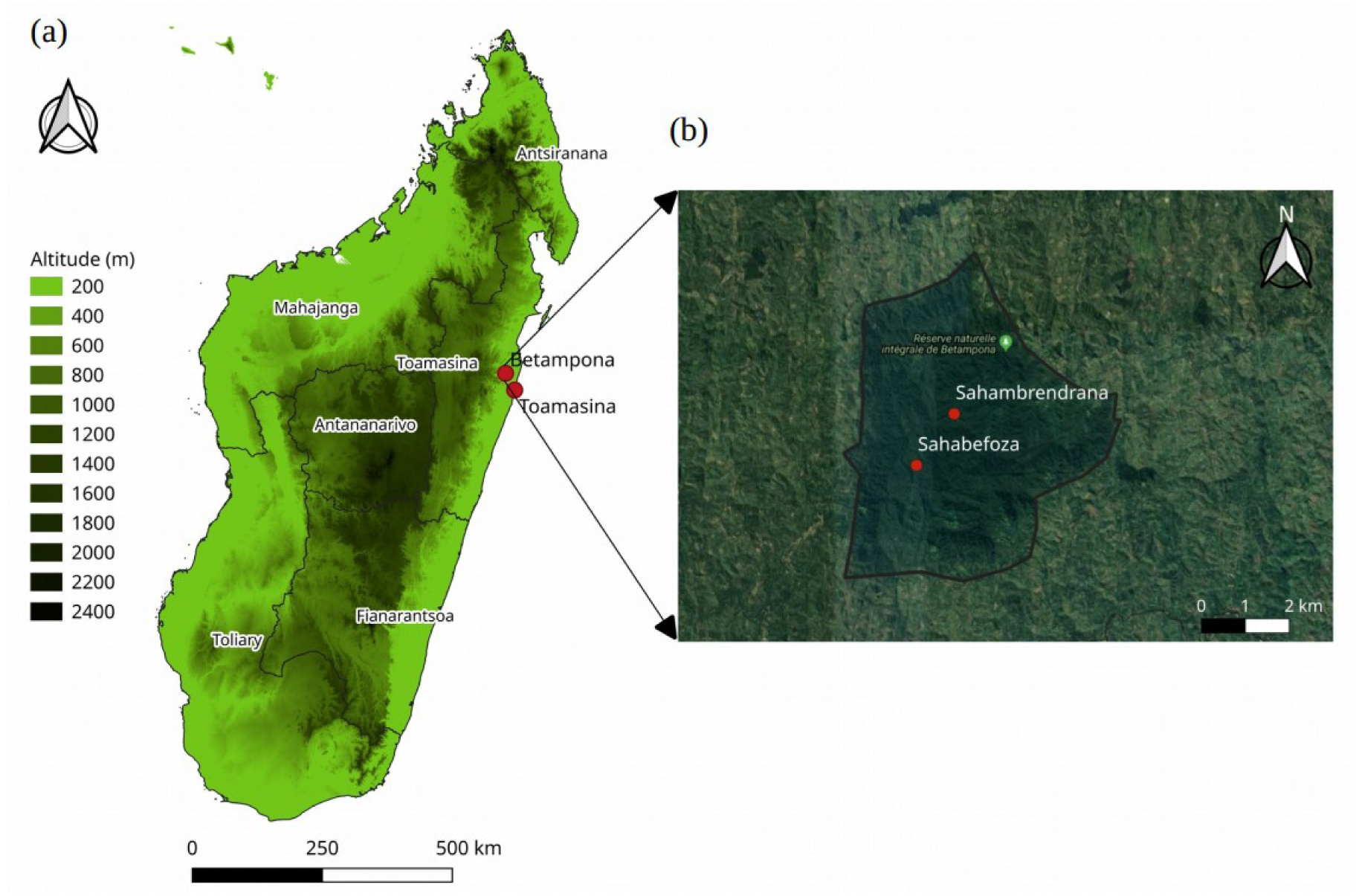
Map representing the relief of Madagascar with the province limits and the localization of Betampona Strict Nature Reserve and Toamasina, the nearest town (raster source: DIVA-GIS) and (b) Betampona forest delimitation with the localization of the two sets of transects monitored.

### II.3 – Data collection and curation

We monitored the activity of amphibian species along transects, either from visual or acoustic detection. At both study sites, we established three transects surveyed year-round between 6:00 pm and 10:00 pm during the 2009-2022 period. In each site, the three transects followed the same scheme: the first one follows a stream, the second one is perpendicular to the first one and follows the slope and the third one is perpendicular to the second one and is along the ridge. The data set contains the presence and abundance of 55 anuran species. These two sites and sets of transects were monitored near-simultaneously (one or two days apart) throughout the study period, but with variable sampling effort between years (Figure 2). We noted the date, site, transect position (along the stream, up slope or on the ridge), species observed and observer identity. We decided to exclude the 2009 data, because in the first year, the fieldworkers were not experienced enough to make reliable species identifications, or the species might have not been detected due to the incompleteness of the data of the year,. We kept all the remaining observations for the community-level analysis (i.e. amphibian total abundance). For the species-level analysis, we removed observation of individuals for which identification was uncertain, or for which the species is yet to be formally described (resulting in 39 species included out of 55). Two species (*Aglyptodactylus inguinalis* and *Boophis roseipalmatus*), were not detected at the beginning of the monitoring, only after 2014 and 2013 (respectively), which is presumably due to the misidentification of those species before these dates rather than their new arrival. Therefore, we removed the data for these species before the corresponding years.

**Figure 2:**
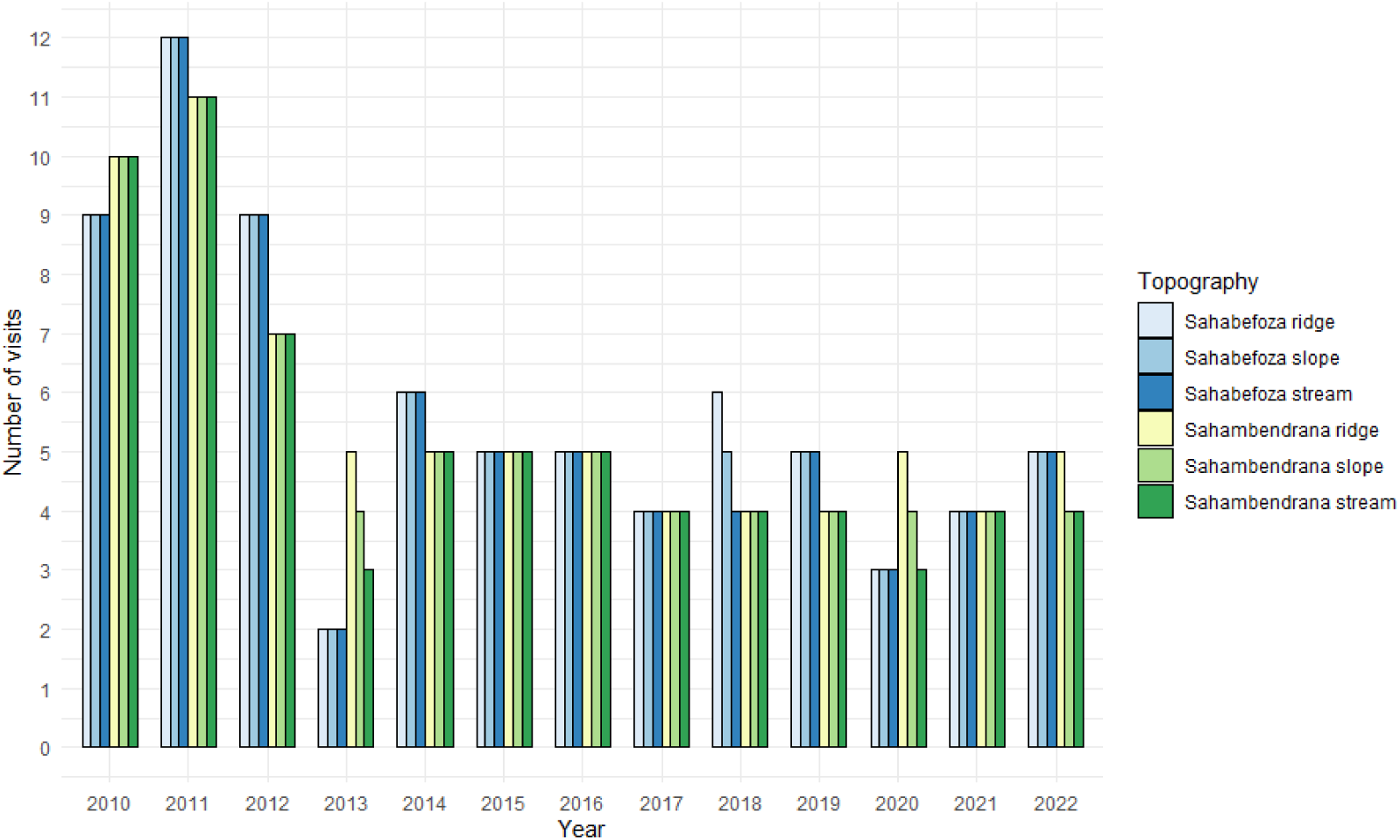
Bar plot representing the heterogeneity in the sampling efforts between years and between the 2 sites (Sahambendrana and Sahabefoza) and the 3 topograpghie (ridge, slope and stream).

### II.2 – Statistical analysis

#### II.2.a – Modelling temporal trends in amphibian total abundance

Amphibians are generally active (and detectable) during specific abiotic conditions. Therefore, detection is more dependent on individual activity than actual abundance. We used activity (i.e. the detection of active individuals) as a proxy for individual count. We cross-tabled the individual counts (individually for each transect to limit zero-inflation) to obtain abundance estimations. We estimated overall linear trends in amphibian activity between 2010 and 2022 based on the coefficients of a linear model with year as a fixed continuous effect. Since amphibian activity is variable within the year (Dubos et al., 2020), estimates of interannual trends may be biased due to our non-homogeneous sampling design. We controlled for non-linear intra-annual (seasonal) variation in species activity using Generalized Additive Mixed Models (GAMMs) Poisson family with Julian date as spline effect (with k fixed to 4, the number of seasons in Madagascar). We retrieved the coefficient of the linear effect of year (i.e. interannual variation) in the presence of the spline effect of Julian date (i.e. intra-annual variation) to control for seasonal fluctuations in activity (Lequy et al., 2017). The Julian date variable has been constructed with the « lubridate » package (Grolemund and Wickham, 2011) of R version 4.4.0 (R Core Team, 2024). The first Julian date was fixed on 1st January. Three other variables were included as random effects, the site, the transect position and the observer group to take into account the differences in detection between observers.

#### II.2.b – Modelling species-specific temporal trends and percentage of change

We selected a subset of species with sufficient observations to estimate trends (> 10 obs.). This approach enabled us to avoid spurious estimations but consequently meant that we could not analyse data for rare species. The 39 species included in this analysis are presented in **Table 1**. Hence, our conclusions are valid for the most common species. We estimated species-specific linear trends using the same modelling approach as formerly described (Part II2a), with the linear coefficient of a year effect from a GAMM (Wood, 2017) adjusting for intra-annual fluctuation in activity, running one model per species. Since some species were observed in a single site or transect, those effects were included in the models as factor effects only when relevant. The observer group was added as a random effect.

**Table 1:** Linear trends of the species for which models have converged. The coefficient (Coef), standard error (Se), and the significance (Si) of the year parameter from the GAMM full models are presented. In the right part of the table, we added the mean and standard deviation (Sd) of the 1,000 coefficients obtained from the resampling bootstrap.

| Species | Complete model |  |  |  | Bootstrap |  |
| --- | --- | --- | --- | --- | --- | --- |
|  | Total | Coef | Se | Si | Mean | Sd |
| *Aglyptodactylus inguinalis*<sup>1</sup> | 35 | 0.0137 | 0.0779 |  | -0.0099<sup>3</sup> | 0.0031 |
| *Blommersia angolafa* | 61 | 0,0341 | 0,0327 |  | 0.0980 | 0.0039 |
| *Boophis albilabris* | 87 | -0,1342 | 0,0365 | *** | -0.0301 | 0.0042 |
| *Boophis albipunctatus*<sup>2</sup> | 12 | -0,2585 | 0,1255 | * | -0,3631 | 0.0237 |
| *Boophis bottae* | 293 | 0,0568 | 0,0178 | ** | 0.0408 | 0.0018 |
| *Boophis madagascariensis* | 1067 | -0,1048 | 0,0095 | *** | -0.0919 | 0.0008 |
| *Boophis roseipalmatus*<sup>1</sup> | 271 | 0.1110 | 0.0247 | ** | 0.1228 | 0.0015 |
| *Boophis rufiocularis* | 571 | -0,0401 | 0,0118 | *** | -0.0365 | 0.0007 |
| *Gephyromantis boulengeri* | 7194 | -0,0369 | 0,0034 | *** | -0.0347 | 0.0004 |
| *Gephyromantis luteus* | 2851 | 0,0014 | 0,0052 |  | -0.0068<sup>3</sup> | 0.0004 |
| *Mantidactylus betsileanus* | 335 | -0,0893 | 0,0167 | *** | -0.0790 | 0.0010 |
| *Mantidactylus lugubris* | 689 | 0,0339 | 0,0100 | *** | 0.0411 | 0.0008 |
| *Platypelis barbouri*<sup>2</sup> | 60 | 0,0775 | 0,0265 | ** | 0.1097 | 0.0122 |
| *Platypelis grandis* | 102 | -0,0368 | 0,0284 |  | -0.0281 | 0.0014 |
| *Platypelis karenae* | 168 | 0,0094 | 0,0203 |  | -0.0065<sup>3</sup> | 0.0018 |
| *Platypelis tuberifera* | 44 | -0,0988 | 0,0441 | * | -0.0872 | 0.0026 |
| *Plethodontohyla notosticta* | 350 | 0,0899 | 0,0137 | *** | 0.0785 | 0.0007 |
| *Stumpffia betampona* | 44 | -0,0335 | 0,0395 |  | -0.0373 | 0.0021 |
| *Stumpffia jeannoeli*<sup>2</sup> | 23 | -0.0832 | 0.0604 |  | -0.1218 | 0.0089 |
| *Anodonthyla* sp. aff. *boulengeri* | 259 | 0,0102 | 0,0169 |  | -0.0019<sup>3</sup> | 0.0009 |
| *Blommersia* sp. Ca13 aff. *grandisonae* | 54 | -0,2159 | 0,0504 | *** | -0.2450 | 0.0038 |
| *Boophis* sp. Ca25 | 312 | 0,0321 | 0,0155 | * | 0.0448 | 0.0013 |
| *Boophis* sp. aff. *luteus* Betampona | 43 | -0,0087 | 0,0401 |  | -0.0243 | 0.0032 |
| *Boophis* sp. aff. *pyrrhus* Sahavontsira | 52 | 0,0860 | 0,0356 | * | 0.0821 | 0.0021 |
| *Boophis* sp. aff. *rappiodes* Betampona<sup>2</sup> | 19 | -0,4146 | 0,1257 | ** | -0.2326 | 0.0198 |
| *Gephyromantis* sp. Ca23 | 455 | -0,1003 | 0,0142 | *** | -0.0904 | 0.0012 |
| *Gephyromantis* sp. aff. *leucomaculatus* | 245 | -0,0066 | 0,0165 |  | -0.0143 | 0.0008 |
| *Gephyromantis* sp. aff. *malagasius* | 2110 | 0,0299 | 0,0059 | *** | 0.0187 | 0.0004 |
| *Gephyromantis* sp. aff. *redimitus* | 2485 | -0,0196 | 0,0056 | *** | -0.0194 | 0.0005 |
| *Guibemantis* sp. aff. *Liber* 1 | 33 | 0,0446 | 0,0486 |  | 0.2314 | 0.0506 |
| *Guibemantis* sp. aff. *punctatus* betampona | 26 | -0,0137 | 0,0516 |  | 0.0239<sup>3</sup> | 0.0054 |
| *Mantidactylus* sp. Ca3 | 1315 | -0,0635 | 0,0081 | *** | -0.0611 | 0.0006 |
| *Mantidactylus* sp. Ca31 | 451 | 0,0157 | 0,0126 |  | 0.0328 | 0.0007 |
| *Mantidactylus* sp. Ca44 | 5027 | -0,0570 | 0,0042 | *** | -0.0521 | 0.0005 |
| *Mantidactylus* sp. Ca55 | 183 | 0,0234 | 0,0192 |  | 0.0154 | 0.0009 |
| *Mantidactylus* sp. Ca56 | 582 | 0,0368 | 0,0109 | *** | 0.0050 | 0.0007 |
| *Mantidactylus* sp. aff. *melanopleura* | 91 | 0,0617 | 0,0268 | * | 0.0392 | 0.0013 |
| *Spinomantis* sp. aff. *aglavei* | 298 | -0,0617 | 0,0172 | *** | -0.0693 | 0.0011 |

<sup>1</sup> For these two species, we removed the first years where they were not recorded. For *Aglyptodactylus inguinalis* we removed 2010 to 2014 data, for *Boophis roseipalmatus* we removed from 2010 to 2013.
<sup>2</sup> For these species, we removed 10% of the coefficient obtained from the bootstrap as it contained aberrant values.
<sup>3</sup> Numbers are in grey when the sign of the mean of the coefficients from bootstrap differs from the sign of the coefficient extracted from full model.
<sup>4</sup> tests are relative to the remaining levels; (\*) $p \leq 0.1$ ; \* $p \leq 0.05$ ; \*\* $p \leq 0.01$ ; \*\*\* $p \leq 0.001$

Once all the estimates were obtained, we predicted the number of individuals estimated by the model at the beginning (2010) and the end (2022) of the monitoring period using the predict function of mgcv package. Then, we computed the percentage change in the number of individuals through time over the 13-year period and used it to propose changes in IUCN status based on two criteria: A2 and B1. Criteria A2 relates to populations with a decrease observed, estimated, inferred or suspected when the cause of the decline may have not ceased nor been understood. Criteria B1 concerns species with small area of occurrence and with a continuing decline observed, estimated, inferred or projected. For Betampona, this latter will be relevant for two micro-endemic species as the total area of Betampona is smaller than 23 km².

#### II.2.c Robustness assessments

##### Robustness of the linear assumptions

In the former approach, we assumed temporal trends in species activity were linear. However, activity may have varied non-linearly throughout the study period, potentially with positive or negative years. To examine interannual variation in activity, we used Generalized Additive Models (GAMs) with year and Julian date as spline effects. We also accounted for site and transect with fixed terms when relevant (i.e. when a given species was observed at multiple sites or transects), and observer group. Then, for each species, we extracted the graphics of the year spline effect for each species.

##### Robustness of sampling design

Since the number of sampling sessions varied greatly between years (ranging between two and 12 per transect; Fig. 2), we assessed the robustness of our models accounting for heterogeneity in the sampling design. For both the community- and species-level approaches, we randomly sampled two dates of sampling sessions by site, transect and year (corresponding to the minimum number of sampling sessions per site/transect/year in our data, i.e., in 2013) and re-ran the same model as described in former sections (II2a and II2b). We repeated the operation 1000 times and compared the distribution of the 1000 ‘year’ effect estimates with the estimate obtained from the full data. We provide the average estimate across the 1000 models. In the species-level approach, we obtained spurious outlier values for a few species with low sample size. We discarded them by removing all estimates that fell into the first or ninth decile (i.e. first or the last 10% of the data) depending on the outlier sign.

## III – Results

From 2009 to 2022, we detected 28,744 amphibians from 55 species (14 genera) over all transects during the 13 years of sampling. All 55 species belong to two families, i.e. Mantellidae (n = 33) and Microhylidae (n = 10), the former being endemic to Madagascar. The number of individuals differed between species, from 1 (*Mantidactylus aerumnalis* and *Scaphiophryne marmorata*) to 7194 (*Gephyromantis boulengeri*). It also differed between transects (ridge: 13,760; slope: 6,575; stream: 8,409), sites (Sahambendrana: 15,202, Sahabefoza: 13,542) and years (from 1,140 in 2020 to 4,909 in 2011). This last difference is due to a heterogeneous effort of sampling between years (from 380 individuals per session and transect in 2020 to 545 in 2011) (**Figure 2**).

### III.2.a – Temporal trends in amphibian total abundance

The amphibian community experienced a significant, but slight decrease during the 13 years of monitoring (*E = −0,0266 +/- 0.0018 ; Pval < 0,001*) (**Figure 3**). On average per site, transect and sampling session, models predicted that 52 individuals (Se +/- 7) were active in 2010 and 38 (Se +/- 5) in 2022, i.e an average loss of 27% of active individuals. The resampling showed a similar trend with a mean estimate of −0.0254 (Sd +/- 0.0002). Predicted values from GAM with year as a spline effect showed 4 trend phases (**Figure 3**). We detected a first important decline from 2010 to 2011, an important decline from 2015 to mid-2017 followed by a strong increase until 2019 then the most important decline until 2022.

**Figure 3:**
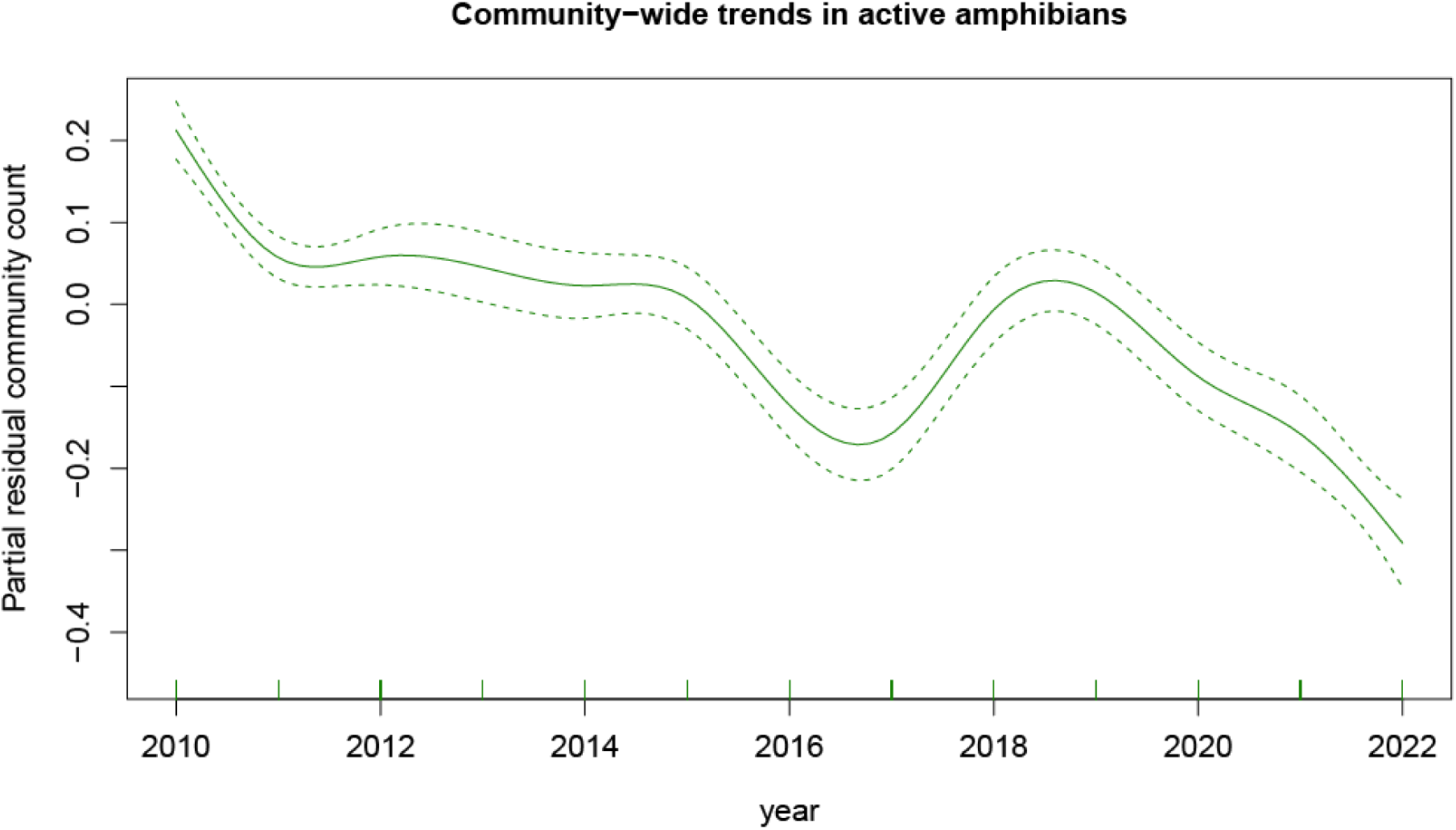
Effect of year on detection of the active amphibians of the community (predicted values obtained from GAMM with the complete model). Dotted lines represent standard errors

### III.2.b – Species-specific temporal trends

Overall, we found a significant increase in activity for 10 species, a significant decrease for 14 species, and no linear trend for 14 species. For decreasing species, the number of active individuals per year decreased by −21% (*Gephyromantis* sp. aff. *redimitus*), down to −99% (*Boophis* sp. aff. *rappiodes Betampona*). For increasing species, the trends ranged from 43% (*Gephyromantis* sp. aff. *malagasius*) to 194% (*Plethodontohyla notosticta*) (**Figure 4**). Predicted values of the spline effect of year showed non-linear variation in the activity of most species, with no consistent pattern between species (**Fig. S1**).

**Figure 4:**
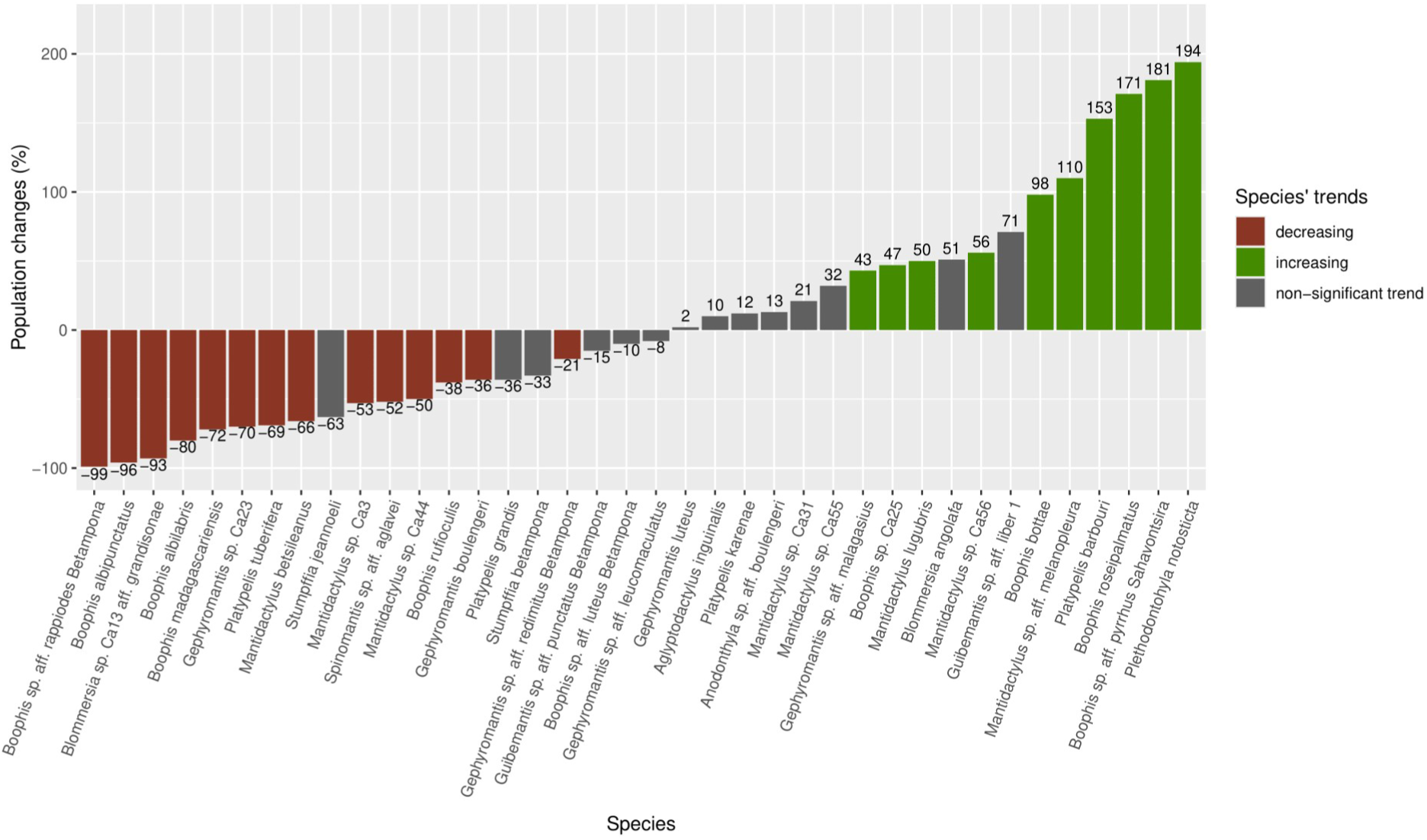
Population trends (expressed as percent change) of 38 species in the amphibian community of Betampona. We computed these changes in abundance of active individuals using model prediction and compare the mean number of individuals per site, transect and sampling session between the beginning and the end of the monitoring period.

The heterogeneous sampling design did not affect our conclusions: in all cases, the direction of the temporal trend was consistent between models calibrated with full data and sub-samples (considering the mean estimate across 1000 iterations), although we found differences in the magnitude of these trends (**Table 1; Fig. S2**).

In the light of these results, we proposed to change species categories according to IUCN criteria A2 and B1 (**Table 2**). Regarding nominal species and following criteria A2, we propose to classify 2 species as Vulnerable (decrease >30%), 2 species as Endangered (decrease >50%) and 2 species as Critically Endangered (decrease >80%). Two other nominal species were assessed considering Criteria B1. They are declining (although not significantly in our models) and are micro-endemic to the Betampona reserve. The size of Betampona is 22.28 km², while the Area of Occurrence (AOO) for a species to be classified as critically endangered is <100 km², provided its decline is at least suspected. We propose then to put these two species in this latter category. Considering candidate species, we propose a classification in a given category, to be applied once they have been formally described. Following criteria A2, 2 species should be classified as critically endangered and 4 as endangered.

**Table 2:** Linear trends of the species from which models showed a significative effect of the year in the full model. We also present the percentage change (% change) of active individuals between the last and first year of the monitoring period, the current IUCN category and a suggestion for an updated one. For two species (*Stumpffia betampona* and *S. jeannoeli*) models didn’t show significative effects but we proposed a change of category based on their trends and micro-endemicity.

| Species | Coef<sup>1</sup> | Si<sup>2</sup> | % change | IUCN Current category | IUCN update proposal |
| --- | --- | --- | --- | --- | --- |
| *Boophis albilabris* | -0,1342 | *** | -80 | LC | CR<sub>A2(b)</sub> |
| *Boophis albipunctatus* | -0,2585 | * | -96 | LC | CR<sub>A2(b)</sub> |
| *Boophis bottae* | 0,0568 | ** | 98 | LC |  |
| *Boophis madagascariensis* | -0,1048 | *** | -72 | LC | EN<sub>A2(b)</sub> |
| *Boophis roseipalmatus* | 0,0824 | ** | 278 | LC |  |
| *Boophis rufiocularis* | -0,0401 | *** | -38 | NT | Vu<sub>A2(b)</sub> |
| *Gephyromantis boulengeri* | -0,0369 | *** | -36 | LC | Vu<sub>A2(b)</sub> |
| *Mantidactylus betsileanus* | -0,0893 | *** | -66 | LC | EN<sub>A2(b)</sub> |
| *Mantidactylus lugubris* | 0,0339 | *** | 50 | LC |  |
| *Platypelis barbouri* | 0,0775 | ** | 153 | LC |  |
| *Platypelis tuberifera* | -0,0988 | * | -69 | LC | EN<sub>A2(b)</sub> |
| *Plethodontohyla notosticta* | 0,0899 | *** | 194 | LC |  |
| *Stumpffia betampona* | -0,0335 |  | -33 | DD | CR<sub>B1(b)</sub> |
| *Stumpffia jeannoeli* | -0,0832 |  | -63 | DD | CR<sub>B1(b)</sub> |
| *Blommersia* sp. Ca13 aff. *grandisonae* | -0,2159 | *** | -93 | NE | CR<sub>A2(b)</sub> |
| *Boophis* sp. Ca25 | 0,0321 | * | 47 | NE |  |
| *Boophis* sp. aff. *pyrrhus* Sahavontsira | 0,0860 | * | 181 | NE |  |
| *Boophis* sp. aff. *rappiodes* Betampona | -0,4146 | ** | -99 | NE | CR<sub>A2(b)</sub> |
| *Gephyromantis* sp. Ca23 | -0,1003 | *** | -70 | NE | EN<sub>A2(b)</sub> |
| *Gephyromantis* sp. aff. *malagasius* | 0,0299 | *** | 43 | NE |  |
| *Gephyromantis* sp. aff. *redimitus* | -0,0196 | *** | -21 | NE |  |
| *Mantidactylus* sp. Ca3 | -0,0635 | *** | -53 | NE | EN<sub>A2(b)</sub> |
| *Mantidactylus* sp. Ca44 | -0,0570 | *** | -50 | NE | EN<sub>A2(b)</sub> |
| *Mantidactylus* sp. Ca56 | 0,0368 | *** | 56 | NE |  |
| *Mantidactylus* sp. aff. *melanopleura* | 0,0617 | * | 110 | NE |  |
| *Spinomantis* sp. aff. *aglavei* | -0,0617 | *** | -52 | NE | EN<sub>A2(b)</sub> |

<sup>1</sup> Coefficient obtained with complete models (ran with full data). Numbers in are in blue and red for species showing a significant positive and negative trend (respectively). For two species, the value of their coefficients is in black because their trends are not significant.
<sup>2</sup> tests are relative to the remaining levels; (\*) $p \leq 0.1$ ; \* $p \leq 0.05$ ; \*\* $p \leq 0.01$ ; \*\*\* $p \leq 0.001$

## IV – Discussion

We assessed amphibian population trends across 13 years in a strict nature reserve from Madagascar with a modelling approach, accounting for heterogeneity in sampling efforts. We found contrasting trends between species, potentially leading to strong changes in future biodiversity patterns and functions. We provide recommendations regarding IUCN conservation status based on criterion A (population change over > 10 years). We suggest updating the status of 15 species towards a threatened category (CR, EN, VU).

### Betampona amphibian trends

Using the only available amphibian long-term monitoring data in the east coast of Madagascar, we studied the population trends of the amphibian community of the Betampona strict nature reserve. We showed that the community has experienced a slight decline in overall abundance since the monitoring began. Species temporal trends differed both in direction and magnitude (15 species declining, 13 species increasing, 13 species with no trend detected).

Comparison of the estimates from the full models and the resampled ones showed very consistent trend directions, although the magnitude differed. This is because resampling produces a very small subset of data, taking two outputs per transect, per site, per year. Count data implies a Poisson distribution assumption for our model (hence a log-transformation of the estimates). The log-link combined with small random samples could explain the large differences in magnitude. We therefore assume, considering the consistent directions of the trends, that our models run with the full data were robust.

In the future, those differential trends will likely change species composition and diversity patterns, and eventually lead to turnover within the community. This explains the slight decrease when we look at the community trend. We choose here to discuss the ecology of nominal species only, as candidate species may have different ecology from their closest counterparts. Of the 5 nominal species showing a positive population trend, 4 are adapted to moderately to highly degraded forests. This is the case of *Mantidactylus lugubris, Boophis roseipalmatus, Plethodontohyla notosticta and Platypelis barbouri* (IUCN SSC Amphibian Specialist Group, 2016a, 2016b, 2016c, 2016d). On the contrary, one increasing species, *Boophis bottae*, is known to live principally in well-preserved forest, even if it can live at the edge of them. Betampona is located at the southern edge of the distribution of some of these species (i.e. *Boophis roseipalmatus* and *Platypelis barbouri*). Given the overall direction of the projected shifts induced by climate change in Madagascar (i.e. upwards and southwards; Raxworthy et al. 2008; Dubos et al. in preprint), climate change may have induced a shift of the optimal conditions for these species. An increase in climate suitability at the level of Betampona induced by climate change may explain these positive trends, which may continue in the future for *Platypelis barbouri*. For *Boophis roseipalmatus*, the population of Betampona is highly isolated and might be exposed to Allee effect (Courchamp et al., 2008), which might counter the increase in the future. For other species such as *Boophis bottae* and *Mantidactylus lugubris*, Betampona is located at the northern edge of their distribution. We fear that their population increase will not endure, and the trends might reverse in the future as a result of climate change.

Surprisingly, many species are in decline despite the high protection status of the Betampona reserve. Only two of them, *Boophis albilabris* and *Boophis rufioculis* are known to live in pristine forest (or only slightly degraded ones). All others can be found in both pristine and degraded areas. Since Betampona is relatively well preserved, these declines might be explained by large-scale processes (e.g. at the landscape level) such as habitat fragmentation which, combined with the low dispersal capacity of amphibians, would drive to the inbreeding of the populations and expose them to the Allee effect. For *Boophis rufioculis*, Betampona represents one highly isolated population, located far from the rest of its distribution (https://www.iucnredlist.org/species/57426/84164900). This suggests that the species is already exposed to both Allee effect and climate change. Further study is needed to disentangle the impact of large- and local-scale processes.

Our results are partially consistent with some studies which have shown a turnover in the species assemblage in forests subject to isolation (Clavel et al., 2011; Schneider-Maunoury et al., 2016). This could be induced by fragmentation, reducing the chance for other populations of specialist species to reach their suitable environments (Andrén et al., 1997; Henle et al., 2004). In our study, generalist species were either in increase or decline. Habitat specialization does not seem to explain the contrasting trends. Further study is needed to better understand the relationship between species ecological traits and their response to global change.

The loss of amphibian abundance can also be the result of other large-scale processes such as climate change, with species being less active or less abundant with changing conditions and resource availability (Dubos et al., 2020). The overall decline may also be due to the isolation of the reserve, which increases the chance of genetic structuring and Allee effect (Courchamp et al., 2008). The decline was not linear, and the factors driving fluctuations throughout the monitoring period remain unclear. The climate of Betampona has already shifted towards warmer and drier conditions (Dubos et al., 2020), and the region is exposed to cyclone activity, which are possible leads to be explored. A study focusing on the drivers of amphibian population trends would help to better understand the underlying mechanisms of amphibian population dynamics and highlight the real threats impacting the Betampona herpetofauna.

### Discussion about the IUCN status

In Madagascar, many species are classified as LC because they are found across a broad geographical range. However, the information on their temporal trends is largely lacking, and some temporal and spatial declines may have occurred, and yet remained undetected. This study highlights the lack of longitudinal data for conservation assessments, leading to underestimation of species extinction risks.

We propose to update the status of 15 species towards a threatened category (CR, EN, VU). For seven of the nominal species for which we propose a change of category, these species are also present in other forests in Madagascar. We justify this choice by the fact that this forest benefits from the highest protection status in Madagascar. We assume that if these species are in decline in this forest, they are likely be in decline in other areas where they are present. Moreover, Betampona is not located at the hot edge of the distributional range of these species, which does not support that they are in decline because they already are reaching the upper edge of their thermal niche (Rehm et al., 2015). For two species (i.e. *Stumpfia betampona* and *S. jeannoeli*, currently classified as Data Deficient), we proposed a status of Critically Endangered despite the estimates of the linear year effects not being statistically significant. As recommended in the IUCN guidelines, we justify our recommendation by the fact that these species are micro-endemic and therefore with an extent of occurrence <100 km², and that we estimated a decline of 33% and 63% over 13 years.

For candidate species, we used the same principle. However, if some species turn out to be micro-endemic after their formal description, their risk of extinction is all the more concerning. Species for which we are proposing a Vulnerable category could therefore be placed in the Critically Endangered category under category B1.

It has been proven that amphibian species with a small geographic range are more susceptible to decline and are more vulnerable to extinction (Chen et al., 2019; Dubos et al., 2022). It is of utmost priority to focus on these species, describe them formally and determine their protection status. This applies to all species classified in the threatened categories of the IUCN, which highlights the need to focus on these species too, including those currently showing a positive trend.

### The threats of forest fragmentation and the importance of protected reserves

The Betampona forest, despite its fully protected status, will nonetheless be subject to a number of threats in the coming years. For example, global changes such as the intensification of cyclones, or the increase in consecutive days of drought will inevitably modify forest and species dynamics (Allan and Liu, 2019; Kumar et al., 2015). Changes in precipitation and humidity will also affect amphibian activity and hence, impact reproduction and population dynamics (Dubos et al., 2020). Light breakthroughs caused by falling trees (due by cyclones or abiotic condition changes inside the forest) will alter water availability, favoring the emergence of invasive exotic species, already present at the forest edge (Subedi et al., 2024; Turton, 1992; Turton and Siegenthaler, 2004). This forest fragment is less than 2,230 ha in size, and a large part of it must or will be subject to the “edge effect”, one of the main consequences of habitat fragmentation. This loss of the buffering capacity of forest fragments is characterized by abiotic changes at the edge of a fragmented forest such as wind speed, increase of solar radiation and temperature (soil and air), and water fluxes (Chen et al., 2019; Laurance et al., 2011; Murcia, 1995; Slater et al., 2024). These changes in environmental conditions are known to severely impact amphibians (Schneider-Maunoury et al., 2016). The magnitude of the edge effect on species depends on the surrounding matrix (Hatfield et al., 2020) and species-specific traits (Ewers and Didham, 2005). One study found an impact of the edge effect up to 1,900 m inside a forest fragment, with a median at 408 m (Schneider-Maunoury et al., 2016). It is worth bearing in mind that the lowest distance between the centroid of Betampona and its frontier is less than 2,000 m, and that there are lots of degraded open zones within the forest. Some authors found that the edge effect can also start from these gaps and spread throughout the forest via a stepping stone phenomenon (Carvajal-Cogollo and Urbina-Cardona, 2015; Sartorius et al., 1999). Given the size of the core forest zone of Betampona, species are very likely to be highly exposed to the edge effect.

Betampona is less than 30 km away from the invasion front zone of the common Asian toad, *Duttaphrynus melanostictus* (Licata, 2022). This toad, known to intoxicate Malagasy snakes (Licata et al., 2022), was first discovered in Madagascar in 2014 (Crottini et al., 2014). It is an explosive breeder, laying around 10,000 eggs per clutch when suitable conditions are met, which contributes greatly to its invasiveness (Deso et al., 2023). Radiotracking studies by the same authors (Licata et al., 2023) have also shown the ability of the invasive toad to disperse both in open and forested environments. This toad will inevitably reach the Betampona forest in the coming years and undoubtedly impact the predators of these fragments, which are used to eating frogs, as already shown in Madagascar for the cat eye snake (*Madagascarophris colubrinus*) (Licata et al., 2022). It will also likely impact native amphibian species, which would be replaced by this highly invasive species.

This study also shows the essential importance of nature reserves. Despite the fact that almost a third of the amphibian species of this forest are in current decline, another third is increasing and another third is represented by species with a flat population trend. One species, *Platypelis karenae*, a Critically Endangered frog endemic to Betampona, shows a stagnating trend, but at least not a declining one. The stagnating trend may indicate that these species are at their population climax for the moment, and have not yet been impacted by global changes. Another possibility (among many) is that they benefit from the availability of new niches induced by other species’ declines and compensate for negative factors. There is a lack of sufficiently solid data to quantify their trends with certainty for the rarest species. In fact, many of these species were rarely seen during monitoring, and we would need to implement dedicated surveys more suited to cryptic species, such as site occupancy during their period of activity.

These results should be seen in the context of the reserve’s highest IUCN protection status. If the decline is so significant in this fragment, one wonders how much more concerning the situation might be in smaller, unprotected forest fragments, which are numerous in Madagascar. However, the protection status in the Madagascar context (of subsistence agriculture) does not guarantee that the habitat will remain unharmed (Piludu et al., 2015). In Betampona, selective logging has already been observed, along with the presence of feral dogs and cats (pers. Obs. Karen Freeman). The arrival of invasive plants, presumably aggravated by the edge effects, represents an additional risk factor. Because the magnitude of edge effect is dependent on the matrix surrounding the fragment, urgent action is needed to mitigate the impact by restoring native trees and controlling invasive alien species at forest edges and in forest gaps. It would also be beneficial, yet hard to implement, to reconnect this fragment of forest, with its high biodiversity value, to the Ankeniheny-Zahamena corridor.

It is more than crucial to continue to develop monitoring in as many fragments as possible, focusing on herpetofauna. With their rapid population dynamics, their high sensitivity to environmental change and their intermediate position within the trophic chain, reptiles and especially amphibians can be useful indicators of the health of an ecosystem and to evaluate the impact of habitat restoration efforts. Some authors recommend a focus on forest core-dependent species as early warning systems (Schneider-Maunoury et al., 2016). Given the problems of knowledge shortfalls and the complexity of the phenomena underlying the global decline of amphibians (Stuart et al., 2004), it is really important to redefine experimental designs that are fit for purpose and focal species. The monitoring of rare or cryptic species must account for their low detectabilit, take into account spatial autocorrelation and probability of detection. This will enable us to understand changes in populations on a fine scale, and to try to isolate for each species the threats that impact them directly.

## V - Conclusions

This study showed a decline in the amphibian community in the Betampona relict forest, a reserve with the highest protection status in Madagascar. The study also revealed contrasting trends between species that might lead to significant changes in amphibian assemblages and functions over time. This study will lay a solid foundation for assessing the status of these different species, for both formally described and candidate species that will be described in the future. Further study is needed to understand the reasons for the declines and the consequences of putative turnovers. It will be important to study the ecology of the different species in greater depth to determine their ecological requirements. We recommend the study potential edge effects in this forest by creating transects running from the forest perimeter towards the center and others running from the gap perimeter towards the forest interior to reveal core-forest zones and set up restoration processes. There is also an urgent need to develop long-term monitoring programs of vertebrates in other forest fragments, often smaller in size that Betampona in the east coast of Madagascar. The development of long-term monitoring programs in the tropics might unveil many unexpected population declines, better identifying species at risk and tackling the decline of biodiversity.

## Notes

### Competing Interest Statement

The authors have declared no competing interest.

